# Molecular mechanism underlying the increased risk of colorectal cancer metastasis caused by single nucleotide polymorphisms in LI-cadherin gene

**DOI:** 10.1101/2022.10.02.510515

**Authors:** Anna Yui, Daisuke Kuroda, Takahiro Maruno, Makoto Nakakido, Satoru Nagatoishi, Susumu Uchiyama, Kouhei Tsumoto

## Abstract

LI-cadherin is a member of the cadherin superfamily. LI-cadherin mediates Ca^2+^-dependent cell-cell adhesion by forming a homodimer. A previous study reported two single nucleotide polymorphisms (SNPs) in the LI-cadherin-coding gene (*CDH17*). These SNPs correspond to the amino acid changes of Lys115 to Glu and Glu739 to Ala. Patients with colorectal cancer carrying these SNPs are reported to have a higher risk of lymph node metastasis than patients without the SNPs. Although proteins associated with metastasis have been identified, the molecular mechanisms underlying the functions of these proteins remain unclear, making it difficult to develop effective strategies to prevent metastasis. In this study, we employed biochemical assays and molecular dynamics (MD) simulations to elucidate the molecular mechanisms by which the amino acid changes caused by SNPs in the LI-cadherin-coding gene increase the risk of cancer metastasis. Cell aggregation assays showed that the amino acid changes weakened the LI-cadherin-dependent cell-cell adhesion. *In vitro* assays demonstrated a decrease in homodimerization tendency due to the mutation of Lys115, and MD simulations suggested an alteration in the intramolecular hydrogen bond network due to the amino acid change. Taken together, our results indicate that the increased risk of lymph node metastasis is due to weakened cell-cell adhesion caused by the decrease in homodimerization tendency.

## Introduction

Liver Intestine-cadherin (LI-cadherin) is a member of the cadherin superfamily. In human body, it is expressed in the normal small intestine and colon cells (1) as well as in various cancer cells, such as gastric adenocarcinoma, colorectal cancer, and pancreatic cancer cells (1–4). In normal intestinal cells, LI-cadherin is located in the intercellular cleft, and the trans-interaction of LI-cadherin is necessary for water transport between the luminal and basal sides (5). Although the role of LI-cadherin in cancer cells has been discussed, the molecular mechanisms by which LI-cadherin contributes to carcinogenesis or tumor progression remain elusive in many cases.

Compared with other members of the cadherin superfamily, LI-cadherin exhibits distinct structural features. LI-cadherin possesses seven extracellular cadherin (EC) repeats, a single transmembrane domain, and a short cytoplasmic domain comprising approximately 20 amino acids (6). Only kidney-specific-cadherin shares structural features with LI-cadherin. Neither of these cadherins belong to any of the previously reported subfamilies (7), and hence, have been termed as 7D-cadherin (8). As is the case with other cadherins that mediate calcium ion-dependent cell–cell adhesion, homodimerization of LI-cadherin is necessary for LI-cadherin-dependent cell–cell adhesion. The unique architecture of the LI-cadherin homodimer and the existence of a noncanonical calcium-free linker, which may have contributed to the formation of this unique antiparallel homodimer, have been shown previously (9).

Aside from LI-cadherin, classical cadherins are well-studied members of the cadherin superfamily. Cadherins, such as E-, N-, and P-cadherins, are classical cadherins. Classical cadherins possess five EC repeats, a single transmembrane domain, and a cytoplasmic domain. Type I classical cadherins exhibit a two-step binding mode. They first form an intermediate “X-dimer” and then a “strand swap-dimer” (ss-dimer) (10–14). Three calcium ions bind to the linker between each EC repeat and contribute to the rigidity of the extracellular region (11). The cytoplasmic domain of classical cadherin consists of more than 100 amino acids and its sequence is conserved within the family. Interaction of this domain with catenins is necessary for efficient cell–cell adhesion via classical cadherins (15, 16). Although LI-cadherin exhibits structural features distinct from classical cadherins, sequence analysis comparing LI-cadherin with E-, N-, and P-cadherins revealed sequence homology between EC1-2 of these proteins as well as between EC3-7 of LI-cadherin and EC1-5 of E-, N-, and P-cadherins (9, 17). The number of EC repeats and the existence of a calcium-free linker in LI-cadherin differentiate both types of cadherins. Another important difference between these cadherins is that the cytoplasmic domain of LI-cadherin does not require interactions with cytoplasmic proteins to maintain cell–cell adhesion. These differences between LI-cadherin and classical cadherins have made it difficult to elucidate the molecular characteristics of LI-cadherin and the underlying mechanisms by which it contributes to cancer progression or metastasis.

A previous study reported two single nucleotide polymorphisms (SNPs) (c.343A>G and c.2216A>C) in the LI-cadherin-coding gene (*CDH17*) (18). These SNPs correspond to the amino acid changes of Lys115 to Glu and Glu739 to Ala, respectively. Patients with colorectal cancer carrying these SNPs are reported to show a higher risk of lymph node metastasis than patients not carrying the SNPs (18). Lymph node metastasis assists the metastasis of cancer cells to distant organs, which decreases the survival rate of patients (19). In line with this, the lymph node ratio has been reported as a promising prognostic indicator for colorectal adenocarcinoma (20). Therefore, it is important to prevent lymph node metastasis in colorectal cancer cells. However, the molecular mechanism by which the risk of lymph node metastasis is increased by these SNPs is not well understood. Understanding the molecular mechanisms of metastasis can aid in the development of therapeutics to prevent cancer metastasis. Considering that both SNPs are responsible for amino acid changes, in this study, we aimed to elucidate how the amino acid changes caused by the SNPs in the LI-cadherin-coding gene increase the risk of cancer metastasis at the molecular level using experimental and computational approaches.

## Results

### Effect of amino acid changes on LI-cadherin-dependent cell–cell adhesion

First, we validated the effects of amino acid changes caused by the SNPs at the cellular level. Considering that cell–cell adhesion mediated by LI-cadherin is maintained by the homodimerization of LI-cadherin (9), cell aggregation assays were employed to compare the cell–cell adhesion ability of the Chinese hamster ovary (CHO) cells expressing WT or mutant LI-cadherin (9, 21, 22). Site-directed mutagenesis was performed to introduce a single or double mutation into the full-length plasmid of EC1-7 fused with monomeric GFP. Single mutations (K115E or E739A) and double mutations (K115E and E739A) were introduced. The established cells were termed K115E-CHO, E739A-CHO, and 2mut-CHO (Figure S1 and Table S1). The cell–cell adhesion abilities of these cells, cells expressing LI-cadherin WT (WT-CHO), and non-transfected mock cells were compared by performing cell aggregation assays (9, 21, 22) and analyzing the size distribution of cell aggregates using micro-flow imaging (MFI).

All cell types expressing LI-cadherin generated cell aggregates. However, the size distribution of cell aggregates was different among the constructs (Figure 1). Cells expressing LI-cadherin mutants generated a larger number of particles than WT-CHO between 25 and 40 µm. In contrast, the number of particles of 40 µm or greater was smaller than that of WT-CHO. The difference in size distribution from that of WT-CHO was more significant when E739A-CHO or 2mut-CHO was used. These results indicate that the mutations caused by the SNPs still allowed the formation of cell aggregates, but at the same time, the mutations inhibited the formation of larger cell aggregates. The Glu739 mutation had a more significant effect than the Lys115 mutation.

**Figure 1.**
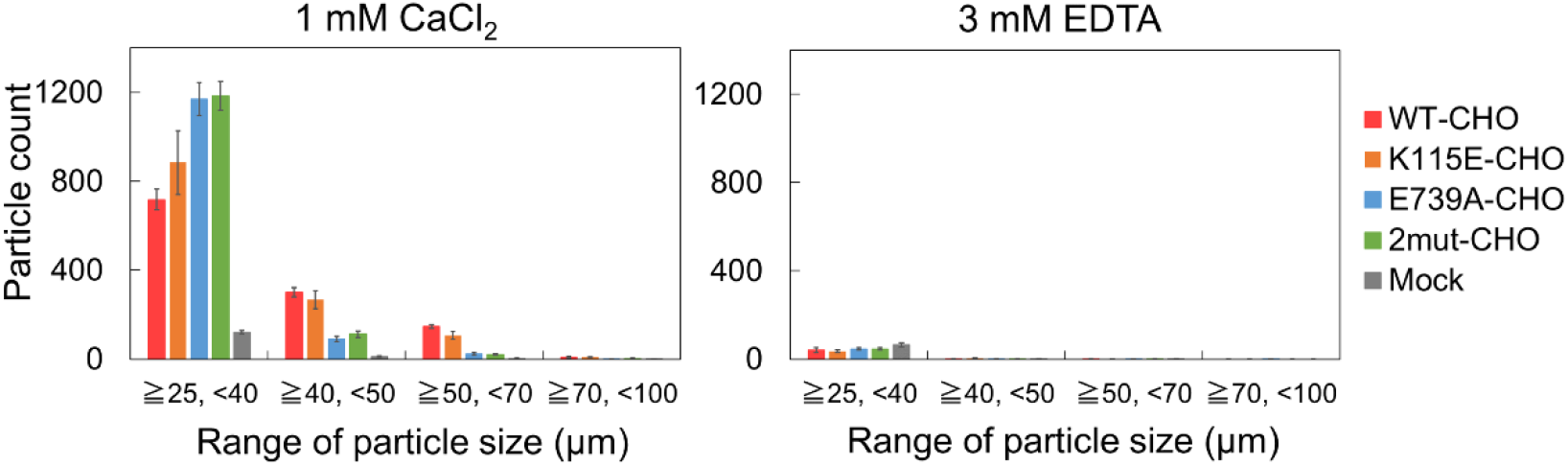
Size distribution of cell aggregates determined via micro-flow imaging (MFI). Particles that are 25 μm or larger were regarded as cell aggregates. LI-cadherin-expressing cells formed cell aggregates when 3 mM EDTA was not added. E739A-CHO and 2mut-CHO tended to form small cell aggregates. Means and standard errors are shown.

We then performed *in vitro* and *in silico* assays to validate the mechanisms underlying decreased cell–cell adhesion ability. Although the mutation of Glu739 had a more significant effect than the mutation of Lys115, we were not able to perform further experimental assays or simulations to investigate the change in molecular characteristics caused by the mutation E739A because the expression level of the recombinant protein containing Glu739, was low, and the crystal structure of the domain containing Glu739, was not available. However, a previous study has suggested that the SNP corresponding to the amino acid change of Lys115 to Glu also has some impact on metastatic potency, depending on the statistical methods employed to analyze the data (18). Therefore, in this study, we focused on biochemical assays and molecular dynamics (MD) simulations of constructs containing Lys115.

### Comparison of homodimerization tendency

Homodimerization of the first four N-terminal domains, EC1-4, is the fundamental step in cell–cell adhesion mediated by LI-cadherin (9). To validate whether the mutation affected the function of LI-cadherin, the homodimerization tendency of WT and K115E of EC1-4 was investigated. Sedimentation velocity-analytical ultracentrifugation (SV-AUC) was used to measure the dissociation constant (*K*_D_) of the homodimer. The *K*_D_ of EC1-4K115E homodimer was 51.6 µM (Figure 2), whereas that of EC1-4WT homodimer was 39.8 µM (9). This result indicates a slight decrease in the homodimerization tendency due to the mutation.

**Figure 2.**
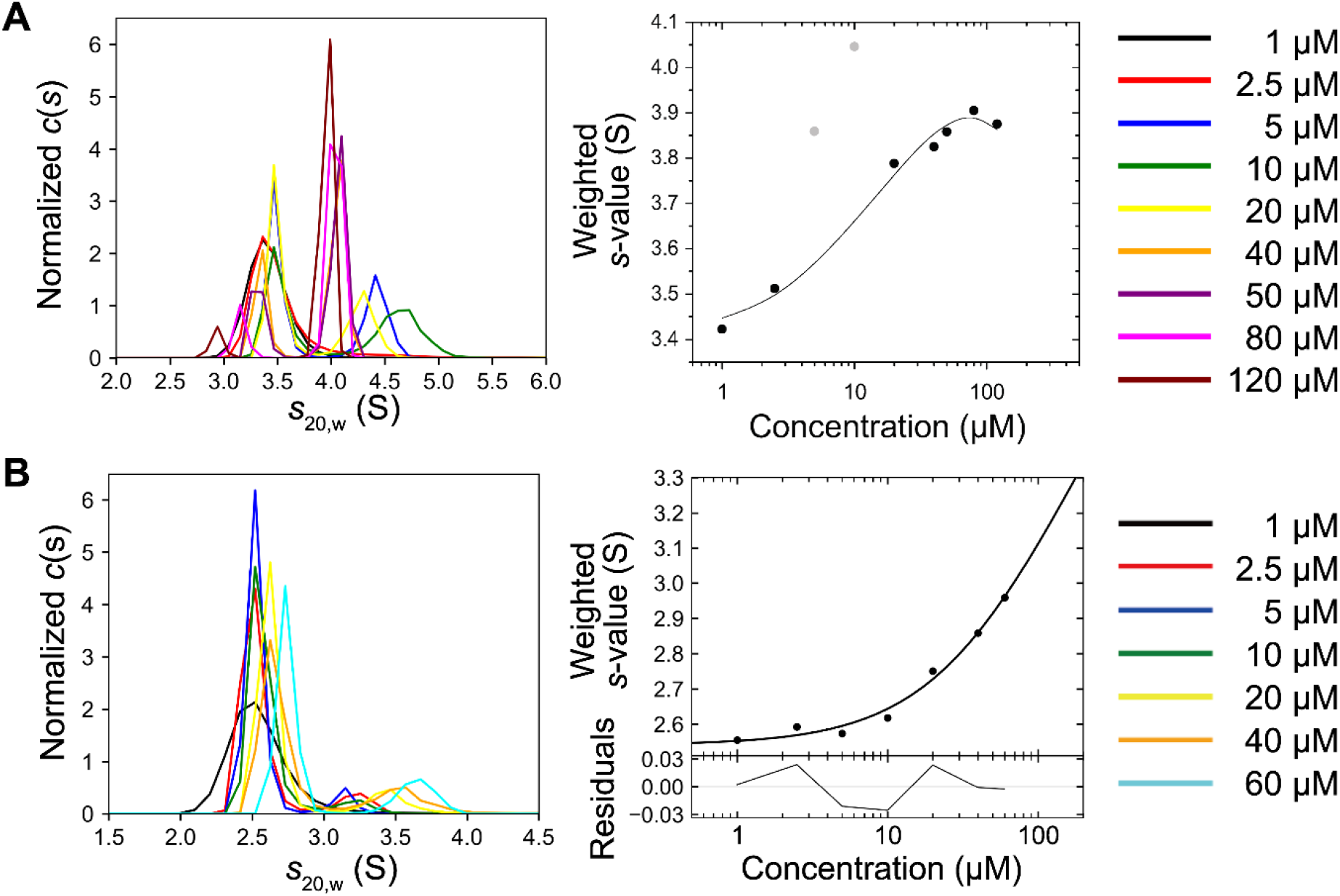
Sedimentation plots of sedimentation velocity-analytical ultracentrifugation (SV-AUC). **A**. EC1-4K115E, **B**. EC1-2K115E. Dimerization of EC1-4K115E and EC1-2K115E was confirmed. *K*_D_ of EC1-4K115E and EC1-2K115E homodimer was 51.6 µM and 343 µM, respectively. *K*_D_ for EC1-2K115E was determined assuming that the *s*-value of EC1-2K115E is the same as that of EC1-2 homodimer which was determined previously (9).

The first two N-terminal domains, EC1-2, also form a homodimer but cannot maintain LI-cadherin-dependent cell–cell adhesion (9). Considering the possibility that the EC1-2 homodimer may be formed in a certain context in the human body, we also performed SV-AUC measurements of the K115E mutant of EC1-2. The *K*_D_ value of the EC1-2K115E was 343 µM, whereas the *K*_D_ value of the EC1-2WT was 75.0 µM (9), showing that the mutation also decreases the homodimerization tendency of EC1-2.

Lys115 is located on the opposite side of the interface of the EC1-4 homodimer and does not directly contribute to the formation of the homodimer (Figure 3). The architecture of the EC1-2 homodimer expressed in *E. coli* has been reported previously (23). The authors pointed out that N-glycans bound to EC2 seemed to hamper the homodimer formation. As EC1-2 used in our experiments was expressed in mammalian cells and N-glycans should be conjugated to EC2 (9), the homodimer architecture formed during our experiment is likely different from the reported structure. Therefore, static crystal structures did not explain how the mutation K115E reduced the ability to form homodimers.

**Figure 3.**
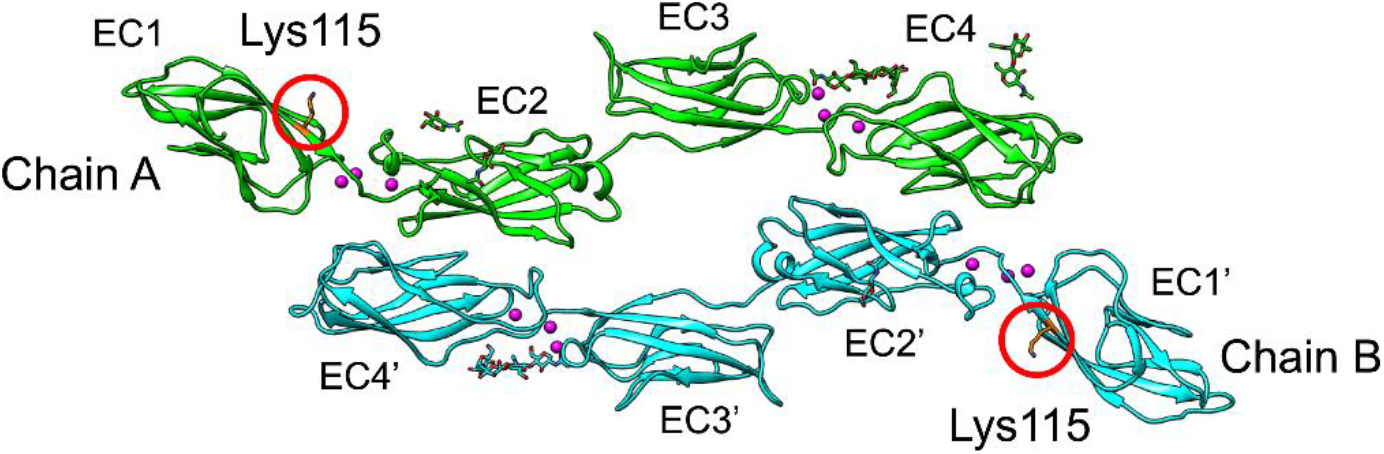
Position of Lys115 mapped on the crystal structure of LI-cadherin EC1-4 homodimer (PDB: 7CYM). Lys115 is shown in orange. Lys115 is located on the opposite side of the homodimer interface.

### Impact of Lys115 to Glu mutation on the structure of LI-cadherin

To explore how the mutation K115E decreased the homodimerization of EC1-4 and EC1-2 at the molecular level, the secondary structures of WT and K115E of EC1-4 and EC1-2 were investigated using circular dichroism (CD) spectroscopy. The CD spectra of all constructs exhibited a large negative peak around 216 nm, which is the typical spectrum of β-sheet-rich proteins (Figure 4A-B), and were consistent with the crystal structure of the EC1-4 homodimer, which showed that EC1, 2, 3, and 4 were β-sheet-rich (9). There was no significant difference between the spectra of the WT and K115E.

**Figure 4.**
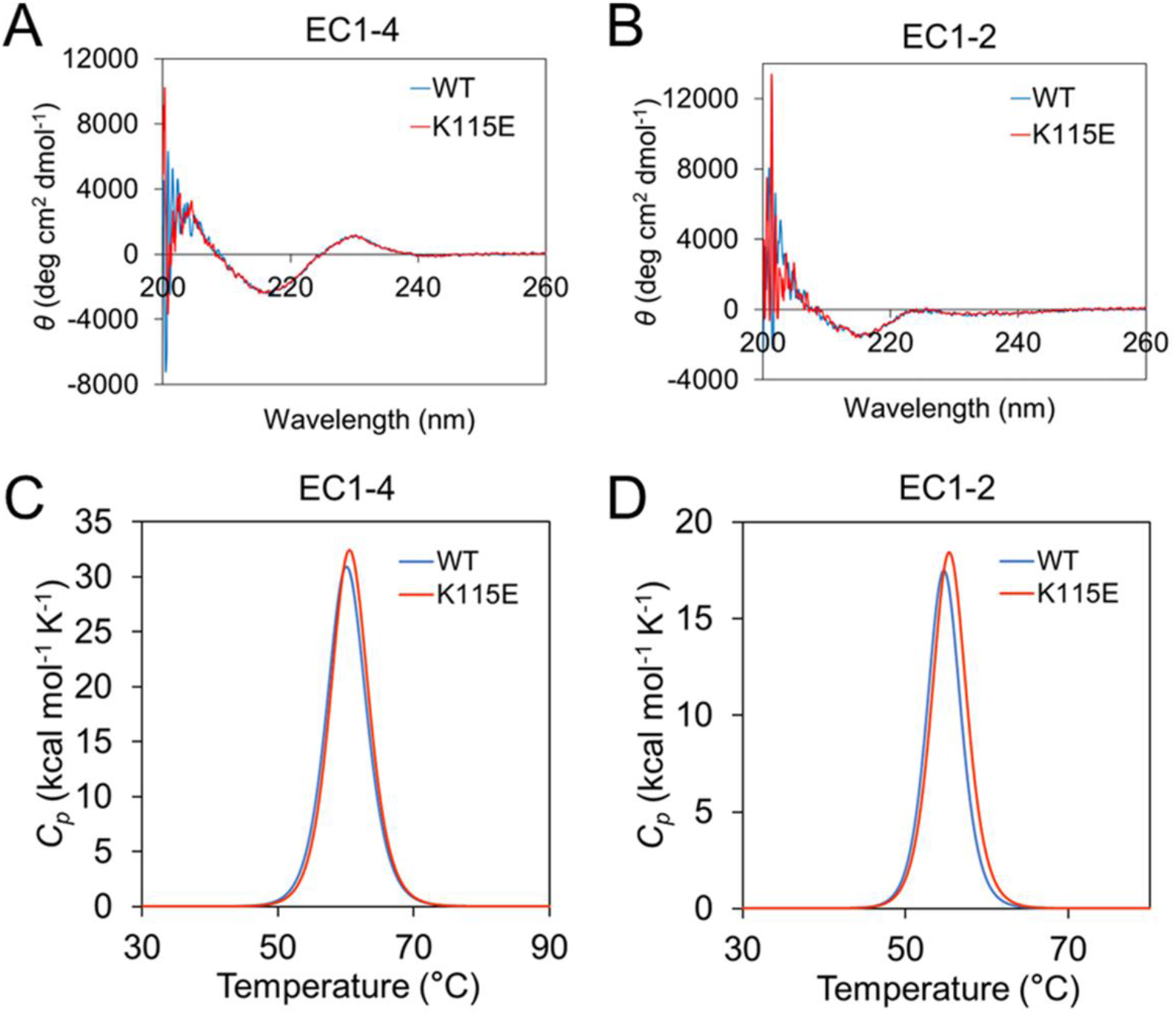
Structural comparison of WT and K115E *in vitro*. **A**. Circular dichroism (CD) spectra of EC1-4WT and EC1-4K115E. **B**. CD spectra of EC1-2WT and EC1-2K115E. **C**. Thermal stability analysis of EC1-4 WT and EC1-4 K115E. **D**. Thermal stability analysis of EC1-2 WT and EC1-2 K115E.

The measurements of thermal stability using differential scanning calorimetry (DSC) also showed that there was no significant difference between *T*_*m*_ and Δ*H* of WT and K115E (Figure 4C-D and Table 1). Both CD and DSC results indicated that the mutation of Lys115 to Glu did not significantly alter the folding of EC1-4 and EC1-2. Therefore, to understand the cause of the decrease in homodimerization tendency caused by the mutation, we next employed MD simulations to investigate the difference between WT and K115E at the atomic level.

**Table 1.** Results of differential scanning calorimetry (DSC) measurements.

| | $T_m$ (°C) | $\Delta H$ (kcal/mol) |
| --- | --- | --- |
| EC1-4WT | 60.1 | 238 |
| EC1-4K115E | 60.5 | 241 |
| EC1-2WT | 54.7 | 95.3 |
| EC1-2K115E | 55.4 | 104 |

### Structural analysis by MD simulations

We performed 400 ns MD simulations of the EC1-4K115E homodimer and compared them with the previous data of the simulations of the EC1-4WT homodimer (Table S1) (9). A mutant structure of Lys115 to Glu in the EC1-4 homodimer was generated using CHARMM-GUI (24, 25). Three independent simulations were conducted at different initial velocities. The convergence of the trajectories was confirmed by calculating the root mean square deviation (RMSD) values of the Cα atoms of each domain, as described in the Experimental Procedures section (Figure S2).

In all simulations, the two chains did not fully dissociate throughout the simulation time. However, when focusing on the dynamics at the interface, clear differences were observed in one of the simulations for the mutant (K115Edimer-Run3). In K115Edimer-Run3, Cα-RMSD of chain B after superposing Cα of chain A indicated that chain B moved drastically with respect to chain A after approximately 100 ns of simulation time while preserving the homodimer structure (Figure 5A-B). Visual inspection of K115Edimer-Run3 showed that part of the interaction observed at the interface of the crystal structure of the homodimer was abolished in the middle of the simulation, whereas the native interactions remained preserved in the simulations of WT and the other two simulations of K115E (Movies S1 and S2). The switching of the hydrophobic interaction partner at the homodimer interface was observed only in K115Edimer-Run3 (Figures S3-S4). This dynamic character at the interface observed in K115Edimer-Run3 may reflect the weaker binding affinity of the homodimer of the mutant than that of the WT, as experimentally measured by SV-AUC.

**Figure 5.**
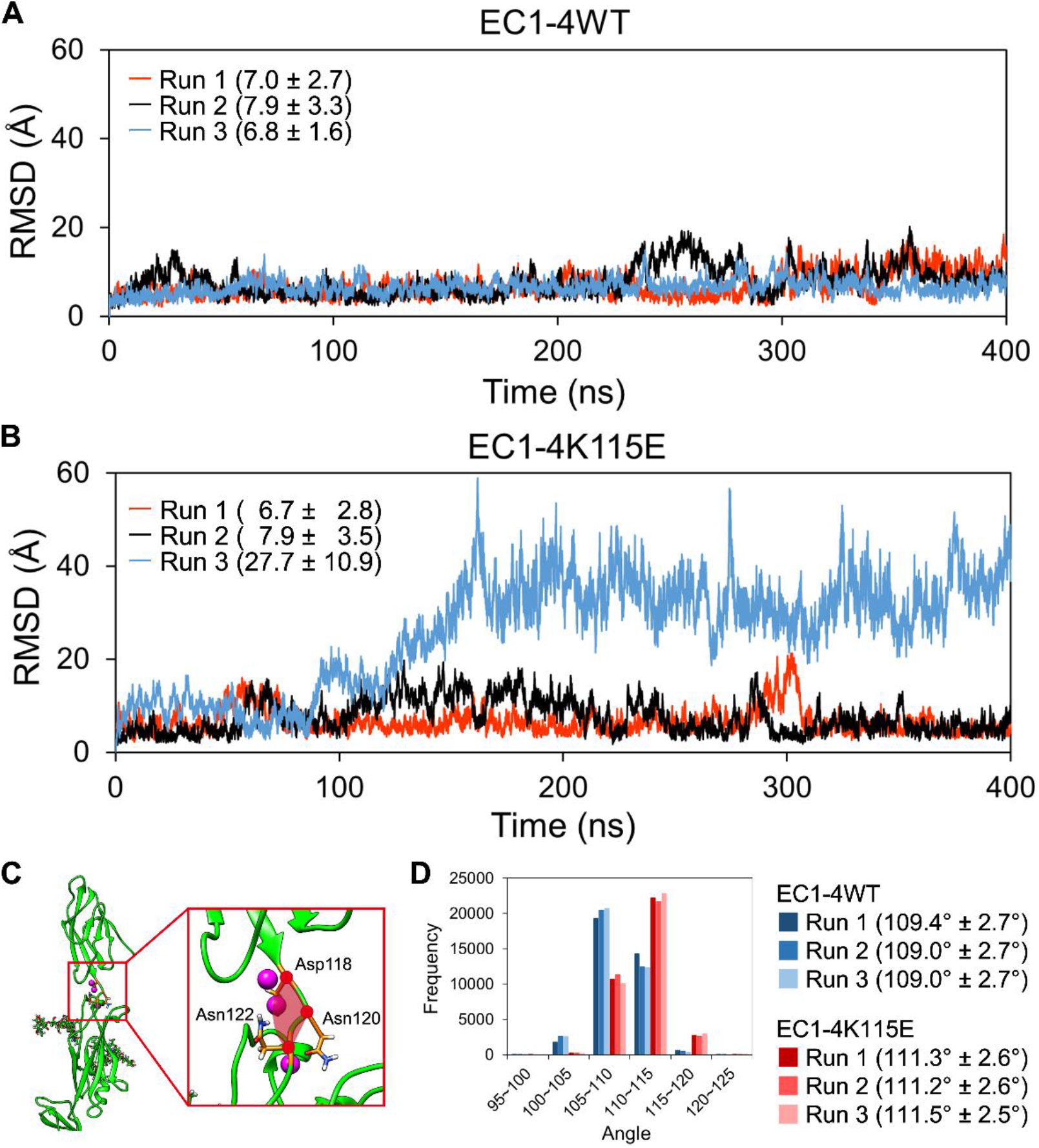
**A**. Root-mean-square deviation (RMSD) values of Cα atoms of chain B after superposing those of chain A in the **A**. EC1-4WT homodimer or **B**. EC1-4K115E homodimer. Averages and standard deviations from 40 to 400 ns of each simulation are shown in the graph in angstrom units. **C**. Angle indicated in red was calculated. **D**. Angle between Asp118, Asn120, and Asn122 was calculated using the GROMACS tool. Averages and standard deviations from 40 to 400 ns of each simulation are shown in parenthesis.

### Causes of differences in the dynamics of the homodimer interface

To further quantify the differences between the WT and the mutant of the EC1-4 homodimer, the angle between EC1 and EC2 was calculated. The angle between the Cα atoms of Asp118, Asn120, and Asn122, which comprise the linker between EC1 and EC2, was defined (Figure 5C). The EC1-4K115E homodimer exhibited a slightly larger angle than that of the EC1-4WT homodimer (Figure 5D).

Next, we investigated the potential source of the change at the interface by analyzing the interaction network around the mutation site (115^th^ residue). First, the distance between the center of gravity of the side chain of the residue at position 115 (Lys115 or Glu115) and Lys117 in chain A of the homodimer was analyzed; the distance was approximately 1.5 Å smaller in K115E (Figure 6). As expected, the distance between OE1/OE2 of Glu115 (Glu115-OE1/OE2) and NZ of Lys117 (Lys117-NZ) in the mutant homodimer was smaller than that between the NZ of Lys115 (Lys115-NZ) and Lys117-NZ in the WT homodimer (Figure S5). This is most likely due to the attractive interaction between negatively charged Glu115-OE1/OE2 and positively charged Lys117-NZ in the mutant and the repulsive interaction between the positively charged Lys115-NZ and Lys117-NZ in the WT. The distance between Glu115-OE1 and Lys117-NZ was ≤ 4 Å in around a quarter of the trajectories of the simulations of the EC1-4K115E homodimer, suggesting the formation of a salt bridge between the two charged residues (Table S2). The formation of hydrogen bonds between the side chains of Glu115 and Lys117 was also observed in simulations of the mutant homodimer (Table 2).

**Table 2.** Percentage of trajectories of which hydrogen bond formation between Lys115 or Glu115 and peripheral residues was confirmed. Trajectories from 40 to 400 ns were considered.

|  |  | Lys117 |  | Thr35~Tyr39 |  | Ser90~Asn93 |  |
| --- | --- | --- | --- | --- | --- | --- | --- |
|  |  | Chain A | Chain B | Chain A | Chain B | Chain A | Chain B |
| WT (%) | Run 1 | 0.0 | 0.0 | 0.0 | 0.2 | 43.1 | 41.3 |
|  | Run 2 | 0.0 | 0.0 | 0.0 | 0.1 | 42.5 | 40.1 |
|  | Run 3 | 0.0 | 0.0 | 0.1 | 0.2 | 47.7 | 39.6 |
| K115E (%) | Run 1 | 39.4 | 37.4 | 4.8 | 6.3 | 5.7 | 5.9 |
|  | Run 2 | 44.5 | 35.1 | 3.7 | 6.5 | 5.6 | 5.8 |
|  | Run 3 | 41.7 | 34.8 | 4.0 | 4.2 | 6.3 | 5.0 |

**Figure 6.**
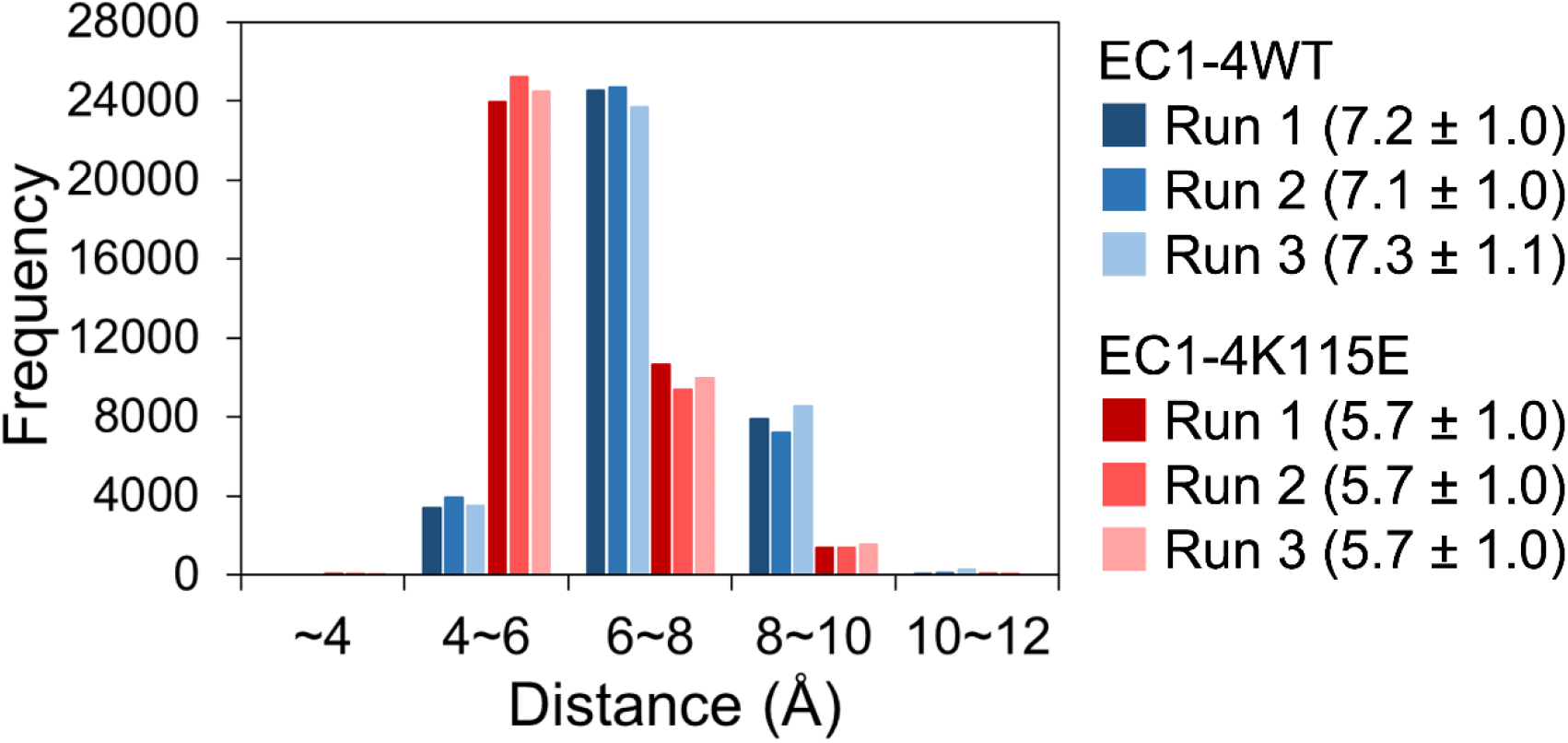
Distance between the center of gravity of Lys115 or Glu115 and Lys117. Means ± standard deviations from 40 to 400 ns in Å units are shown.

The hydrogen bond network between the 115^th^ residue and residues in the adjacent β-strands was also investigated. The number of hydrogen bonds between the 115^th^ residue and residues from the 35^th^ to 39^th^ or from the 90^th^ to 93^rd^ positions was analyzed. The frequency of hydrogen bond formation between the 115^th^ residue and residues from 35^th^ to 39^th^ positions increased in the mutant homodimer (Table 2). In contrast, the number of hydrogen bonds between the 115^th^ residue and residues from the 90^th^ to 93^rd^ positions was significantly decreased by the K115E mutation (Table 2). These changes in the hydrogen bond network appear to have contributed to the difference in the angle between EC1 and EC2, and may explain the slight difference in *T*_*m*_ and Δ*H* of WT and K115E observed in the DSC measurement.

Although a clear alteration of the interface dynamics occurred only in K115Edimer-Run3, changes in the hydrogen bond network were observed in two other runs of K115E. These results suggest that the mutation of Lys115 could significantly alter the dynamics of the molecule by slightly adjusting the folding of the molecule.

## Discussion

Metastasis is a major factor that increases cancer mortality. Understanding the mechanisms underlying cancer metastasis is essential for developing effective cancer treatment strategies. Although many types of proteins have been identified as key factors that govern cancer metastasis, only a few studies have revealed the mechanism underlying metastasis by focusing on the molecular characteristics of target proteins. Here, we have shown the possible roles of LI-cadherin in lymph node metastasis of colorectal cancer by analyzing how amino acid changes caused by SNPs in the LI-cadherin-coding gene affect cell aggregation ability and the conformation of the molecule.

Our data showed that the mutations K115E and E739A decreased cell aggregation. *In vitro* and *in silico* analyses have shown that the mutation K115E decreases homodimerization tendency, possibly due to changes in the hydrogen bond network around the 115^th^ residue. Collectively, as a mechanism of how SNPs in the LI-cadherin gene increase the risk of colorectal cancer metastasis, we propose the mechanism shown in Figure 7. As LI-cadherin-dependent cell–cell adhesion is achieved by homodimerization of LI-cadherin, cell–cell adhesion is weakened by a decrease in homodimerization tendency. The weakened cell–cell adhesion ability leads to a higher risk of cancer cell migration from the primary tumor, which leads to a higher risk of metastasis.

**Figure 7.**
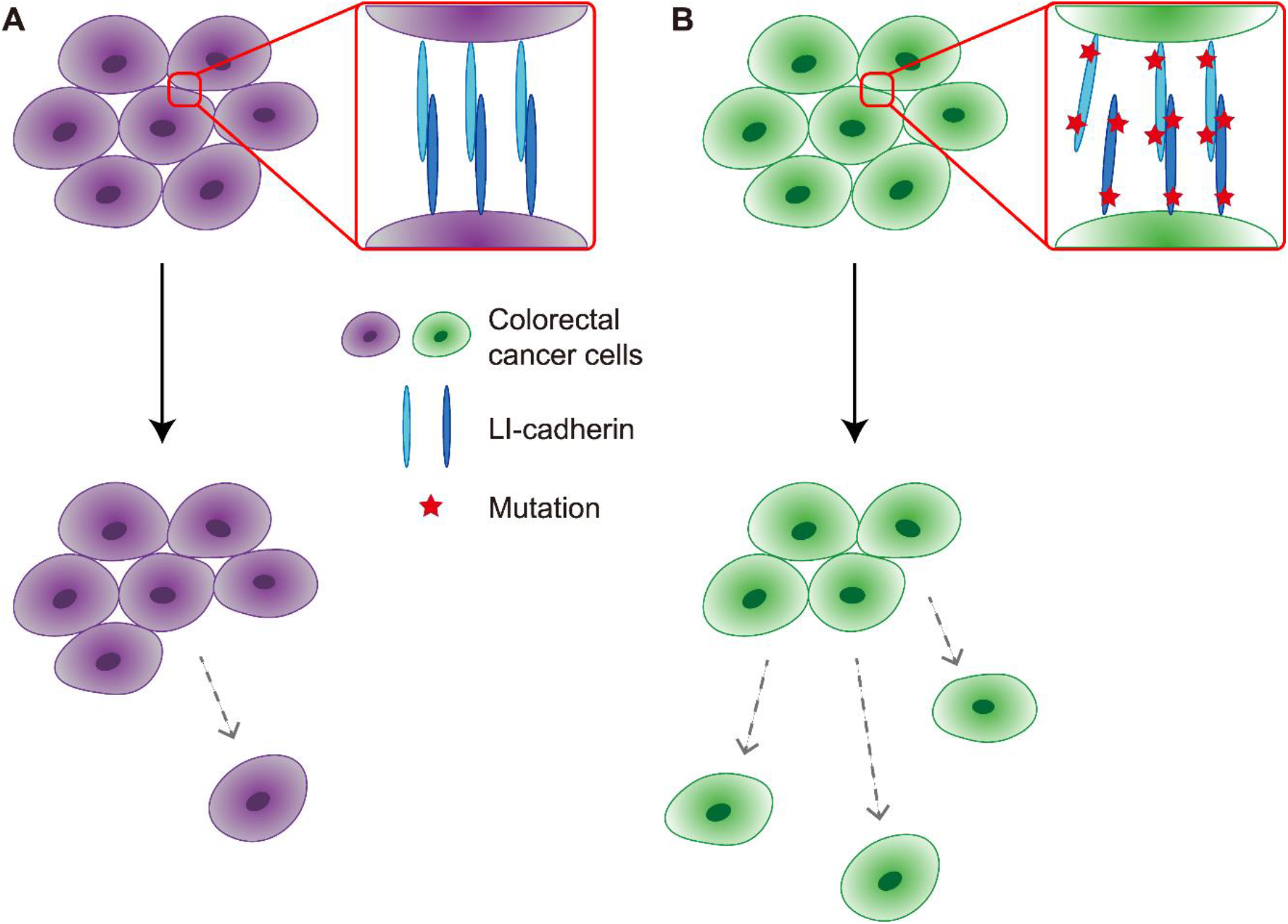
Possible mechanisms of increased lymph node metastasis of colorectal cancer associated with single nucleotide polymorphisms (SNPs) in LI-cadherin-coding gene. **A**. Patients without SNPs. **B**. Patients with SNPs.

A previous study on the genomic analysis of colorectal cancer patients has shown that the G/G genotype or G allele of c.343 and C/C genotype or C allele of c.2216 increase the risk of lymph node metastasis. However, when another statistical method was employed, no difference in lymph node metastasis was observed between the G allele and A allele of c.343, whereas the difference was still observed between the C allele and A allele of c.2216 (18). Our cell aggregation assay data showed that the Glu739 mutation weakened cell aggregation ability more than the Lys115 mutation, suggesting that the impact of the Lys115 mutation is also smaller in the human body. We speculated that our *in vitro* and *in silico* assays were able to capture subtle effects induced by the mutation, which may reflect the results of a previous study showing that the results of the statistical assessment differed between the methods employed.

The contribution of LI-cadherin to cancer progression depends on the tumor type. A study using the human colorectal adenocarcinoma cell line LoVo showed that knockdown of LI-cadherin increased the invasion and metastatic potency of LoVo cells (26). In contrast, knockdown of the LI-cadherin gene in the mouse pancreatic ductal adenocarcinoma cell line Panc02-H7 suppressed the cell proliferation *in vitro* and orthotopic tumor growth *in vivo* (4). This was also the case for E-cadherin expression. Whereas loss of cell–cell adhesion ability by the downregulation of E-cadherin by epithelial–mesenchymal transition is often observed in cancer cells (27, 28), it has also been reported that E-cadherin promotes metastasis in various invasive ductal carcinoma models by limiting apoptosis mediated by reactive oxygen species (29). Our results suggest that in cancer cells in which LI-cadherin suppresses metastasis, molecules that strengthen LI-cadherin-dependent cell–cell adhesion can be used to suppress the migration of cancer cells by maintaining LI-cadherin-dependent cell–cell adhesion.

To the best of our knowledge, this is the first report to describe the structural basis for the increased risk of cancer metastasis caused by amino acid changes in cell–cell adhesion molecules. Cancer metastasis is a complex process involving the invasion of primary tumor, its intravasation into blood vessels and lymph nodes, circulation within blood vessels and lymph nodes, extravasation, and settlement at the metastatic site (30). In the lymph node metastasis of colorectal cancer focused on in this study, we believe that not only LI-cadherin-dependent cell–cell adhesion, but also various other factors, such as interaction between LI-cadherin and other cell surface proteins or extracellular matrix components and intracellular signal transduction, may contribute. Although further studies, including the use of *in vivo* models, are needed to fully describe the mechanisms by which SNPs on the LI-cadherin gene increase the risk of lymph node metastasis in colorectal cancer, our study highlighted the contribution of cell– cell adhesion molecules to the complex interaction network of cancer metastasis, suggesting that molecules targeting cell–cell adhesion proteins may have the potential to inhibit cancer metastasis.

### Experimental procedures

#### Expression and purification of recombinant LI-cadherin

All LI-cadherin constructs were expressed as previously described (9). Briefly, Expi293F cells (Thermo Fisher Scientific) were transfected with plasmids containing the LI-cadherin sequence, following the manufacturer’s protocol. Cells were harvested at 37 °C and 8.0% CO_2_ for three days after transfection. The supernatant was dialyzed against a solution of 20 mM Tris-HCl (pH 8.0), 300 mM NaCl, and 3 mM CaCl_2_. Proteins were purified by immobilized metal affinity chromatography using Ni-NTA Agarose (Qiagen), followed by size exclusion chromatography using the HiLoad 26/600 Superdex 200 pg column (Cytiva) at 4 °C equilibrated in buffer A (10 mM HEPES-NaOH (pH 7.5), 150 mM NaCl, and 3 mM CaCl_2_). Unless otherwise indicated, samples were dialyzed in buffer A before analysis, and the filtered dialysis buffer was used for assays.

#### Site-directed mutagenesis

Introduction of mutations in plasmids was performed as described previously (31).

#### Circular dichroism (CD) spectroscopy

CD spectra were measured using a J-820 spectrometer (JASCO) at 25 °C. Spectra from 260 to 200 nm were scanned five times at a scan rate of 20 nm/min. A quartz cuvette, with an optical length of 1 mm, was used. The measurements were performed at protein concentrations of 5 µM (EC1-4WT and EC1-4K115E) and 10 µM (EC1-2WT and EC1-2K115E). The spectrum of the buffer was subtracted from that of each protein sample. Data are represented as mean residue ellipticities by normalizing the subtracted spectra with protein concentration and the number of amino acid residues comprising the protein.

#### Differential scanning calorimetry (DSC)

DSC measurements were performed using a MicroCal PEAQ-DSC Automated System (Malvern Panalytical). The measurements were performed from 10 to 100 °C at a scan rate of 1 °C/min. The protein concentrations were 20 µM (EC1-4WT and EC1-4K115E) and 30 µM (EC1-2WT and EC1-2K115E). Data were analyzed using the MicroCal PEAQ-DSC software.

#### MD simulations

MD simulations of LI-cadherin were performed as previously described (9). In brief, GROMACS 2016.3 (32) with a CHARMM36m force field (33) was used. The crystal structure of the EC1-4WT homodimer (PDB: 7CYM) was used as the initial structure. The sugar chains were removed from the original crystal structure. Missing residues were modelled using MODELLER 9.18 (34). The addition of N-linked sugar chains (G0F) and the mutation of Lys115 to Glu115 was performed using CHARMM-GUI (24, 25). The structures were solvated with TIP3P water (35) in a rectangular box such that the minimum distance to the edge of the box was 15 Å under periodic boundary conditions using CHARMM-GUI (24). Each system was energy-minimized for 5000 steps and equilibrated with the NVT ensemble (298 K) for 1 ns. Further simulations were performed using the NPT ensemble at 298 K. The time step was set as 2 fs throughout the simulation. A snapshot was recorded every 10 ps. All trajectories were analyzed using the GROMACS tools.

#### Sedimentation velocity analytical ultracentrifugation (SV-AUC)

The detailed protocol for SV-AUC has been described in a previous report (9). Briefly, the Optima AUC (Beckman Coulter) equipped with an 8-hole An50 Ti rotor was used. Measurements were performed at 20 °C with 1, 2.5, 5, 10, 20, 40, 50, 80, and 120 µM of EC1-4K115E or 1, 2.5, 5, 10, 20, 40, and 60 µM of EC1-2K115E. A protein sample (390 µL) was loaded into the sample sector of a cell equipped with sapphire windows and 12 mm double-sector charcoal-filled on the centerpiece. Buffer (400 µL) was loaded into the reference sector of each cell. Data were collected at 42,000 rpm with a radial increment of 10 µm, using a UV detection system.

#### Establishment of CHO cells expressing LI-cadherin mutants

CHO cells expressing LI-cadherin mutants were established as described previously (9). The DNA sequence of the LI-cadherin constructs fused with monomeric GFP was cloned into the pcDNA5/FRT vector (Thermo Fisher Scientific). The Flp-In-CHO Cell Line (Thermo Fisher Scientific) was transfected with the plasmid, following the manufacturer’s protocol. Cloning was performed using the limiting dilution-culture method. Cells were observed using an In Cell Analyzer 2000 instrument (Cytiva) with the FITC filter (490/20 excitation, 525/36 emission), and the cells expressing GFP were selected. The cells were cultivated in Ham’s F-12 Nutrient Mixture (ThermoFisher Scientific) supplemented with 10 % fetal bovine serum (FBS), 1% L-glutamine, 1% GlutaMAX-I (ThermoFisher Scientific), 1% penicillin–streptomycin, and 0.5 mg mL^-1^ hygromycin B at 37 °C and 5.0% CO_2_.

#### Cell imaging

After establishing the cell lines expressing LI-cadherin, the expression of LI-cadherin in cells was further confirmed by observing the fluorescence of GFP. The protocols for imaging the cells have been previously described (9).

#### Cell aggregation assay

The cell aggregation assays were performed by modifying the methods described in previous reports (21, 22). The detailed protocol has been previously described (9). Briefly, LI-cadherin expressing cells were detached from the plate using 0.01% trypsin and washed with 1x HMF supplemented with 20% FBS. Cells were suspended in 1x HMF and 5 × 10^4^ cells were loaded into a 24-well plate coated with 1% w/v BSA. EDTA was added if necessary. After incubating the plate at room temperature for 5 minutes, 24-well plate was placed on a shaker and rotated at 80 rpm for 60 minutes at 37 °C.

#### Micro-flow Imaging (MFI)

MFI (Brightwell Technologies) was used to count the particle number and visualize the cell aggregates after the cell aggregation assays as described previously (9). After the cell aggregation assay procedure described above, the plate was incubated at room temperature for 10 minutes, followed by the addition of 500 µL of 4%-Paraformaldehyde Phosphate Buffer Solution (Nacalai Tesque) to each well. The plate was then incubated on ice for more than 20 minutes. The MFI View System Software was used for measurements and analyses. The instrument was flushed with detergent and ultrapure water before the experiments. The cleanliness of the flow channel was checked by performing measurements using ultrapure water and confirming the presence of less than 100 particles/mL. The flow path was washed with 1x HMF before the samples were measured. The purge volume and analyzed volume were 200 and 420 µL, respectively. Optimize Illumination was performed prior to each measurement. The number of particles that were 1 µm or larger and less than 100 µm in size was counted.

## Acknowledgments

We thank Dr O. Kusano-Arai, Dr H. Iwanari, and Dr T. Hamakubo for providing us with gene sequence of LI-cadherin. The supercomputing resources in this study were provided by the Human Genome Center at the Institute of Medical Science, The University of Tokyo, Japan.

## Funding and additional information

This work was funded by a Grant-in-Aid for Scientific Research (A) 16H02420 (to K. T.) and a Grant-in-Aid for Scientific Research (B) 20H02531 (to K. T.) from the Japan Society for the Promotion of Science, a Grant-in-Aid for Scientific Research on Innovative Areas 19H05760 and 19H05766 (to K. T.) from the Ministry of Education, Culture, Sports, Science and Technology, and a Grant-in-Aid for JSPS Fellows 18J22330 (to A. Y.) from the Japan Society for the Promotion of Science.

