## Supporting Information for "Molecular mechanism underlying the increased risk of colorectal cancer metastasis caused by single nucleotide polymorphisms in LI-cadherin gene"

**Running title**: How SNPs in LI-cadherin gene increase metastasis

**Table S1: Constructs of LI-cadherin used in this study.**
**Table S2: Percentage of trajectories from 40 to 400 ns of which the distance between Lys115-NZ or Glu115-OE1 and Lys117-NZ was less than 4 Å.**

**Figure S1: Verification of LI-cadherin expression in CHO cells.**
**Figure S2: RMSD values of Cα atoms during the simulations of EC1-4K115E homodimer.**
**Figure S3: Distance between the center of gravity of Ile169 or Leu171 in chain A and Ala373 or Phe376 or Ala400 in chain B during the simulations of the A. EC1-4WT homodimer or B. EC1-4K115E homodimer.**
**Figure S4: Distance between the center of gravity of Met173 in chain A and Ala400 or Met402 in chain B during the simulations of the A. EC1-4WT homodimer or B. EC1-4K115E homodimer.**
**Figure S5: Distance between the NZ atom of Lys117 and A. NZ atom of Lys115 in the EC1-4WT homodimer, B. OE1 atom, or C. OE2 atom of Glu115 in the EC1-4K115E homodimer.**

**Movie S1. K115Edimer-Run3**
**Movie S2. WTdimer-Run1**

**Table S1: Constructs of LI-cadherin used in this study.**

| Construct name | Sequence[c] | Usage |
|---|---|---|
| EC1-4WT[a]<br>EC1-4K115E[a] | 23–441 | CD and DSC |
| EC1-2WT[a]<br>EC1-2K115E[a] | 23–236 | CD and DSC |
| WT-CHO[b]<br>K115E-CHO[b]<br>E739A-CHO[b]<br>2mut-CHO[b] | 23–832 | Cell aggregation assays |
| EC1-4WT<br>EC1-4K115E | 23–441 | MD simulations |

a. Myc-tag (EQKLISEEDL), NSAVD sequence, and His-tag (HHHHHH) were added to the C-termini of these sequences.

b. Gly-Ser linker and monomeric GFP were added to the C-terminus.

c: Residue numbering follows the numbering of Uniprot entry Q12864.

**Table S2. Percentage of trajectories from 40 to 400 ns of which the distance between Lys115-NZ or Glu115-OE1 and Lys117-NZ was 4 Å or less.**

| | WT | | | | K115E | | |
|---|---|---|---|---|---|---|---|
| | Run 1 | Run 2 | Run 3 | | Run 1 | Run 2 | Run 3 |
| Lys115-NZ (%) | 0.00833 | 0.0222 | 0.00278 | Glu115-OE1 (%) | 25.0 | 27.5 | 26.0 |
| | | | | Glu115-OE2 (%) | 24.3 | 29.0 | 26.2 |

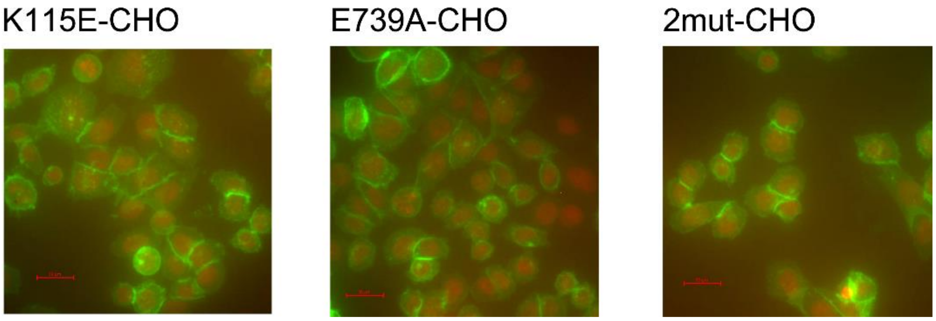


**Figure S1.** Verification of LI-cadherin expression in CHO cells. Expression of LI-cadherin on the cell surface was confirmed by observing the fluorescence of GFP with an In Cell Analyzer 2000 instrument. Fluorescence of GFP and Hoechst 33342 are indicated in green and red, respectively. Expression of LI-cadherin was observed in all constructs shown above. Scale bars represent 20 μm. LI-cadherin expression on the cell surface of WT-CHO was previously confirmed (9).

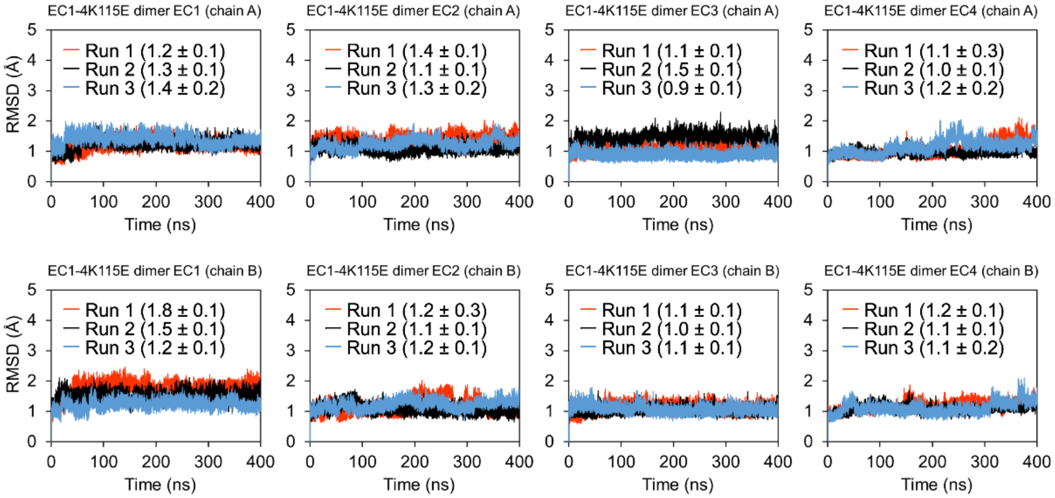


**Figure S2.** RMSD values of Cα atoms during the simulations of EC1-4K115E homodimer. As the molecule showed high flexibility at $Ca^{2+}$-free linker, RMSD values of each domain was calculated individually. Five Cα atoms at N-terminus were excluded from the calculation of RMSD of EC1 as they were disordered. Averages and standard deviations from 40 to 400 ns are shown in the graph in angstrom unit.

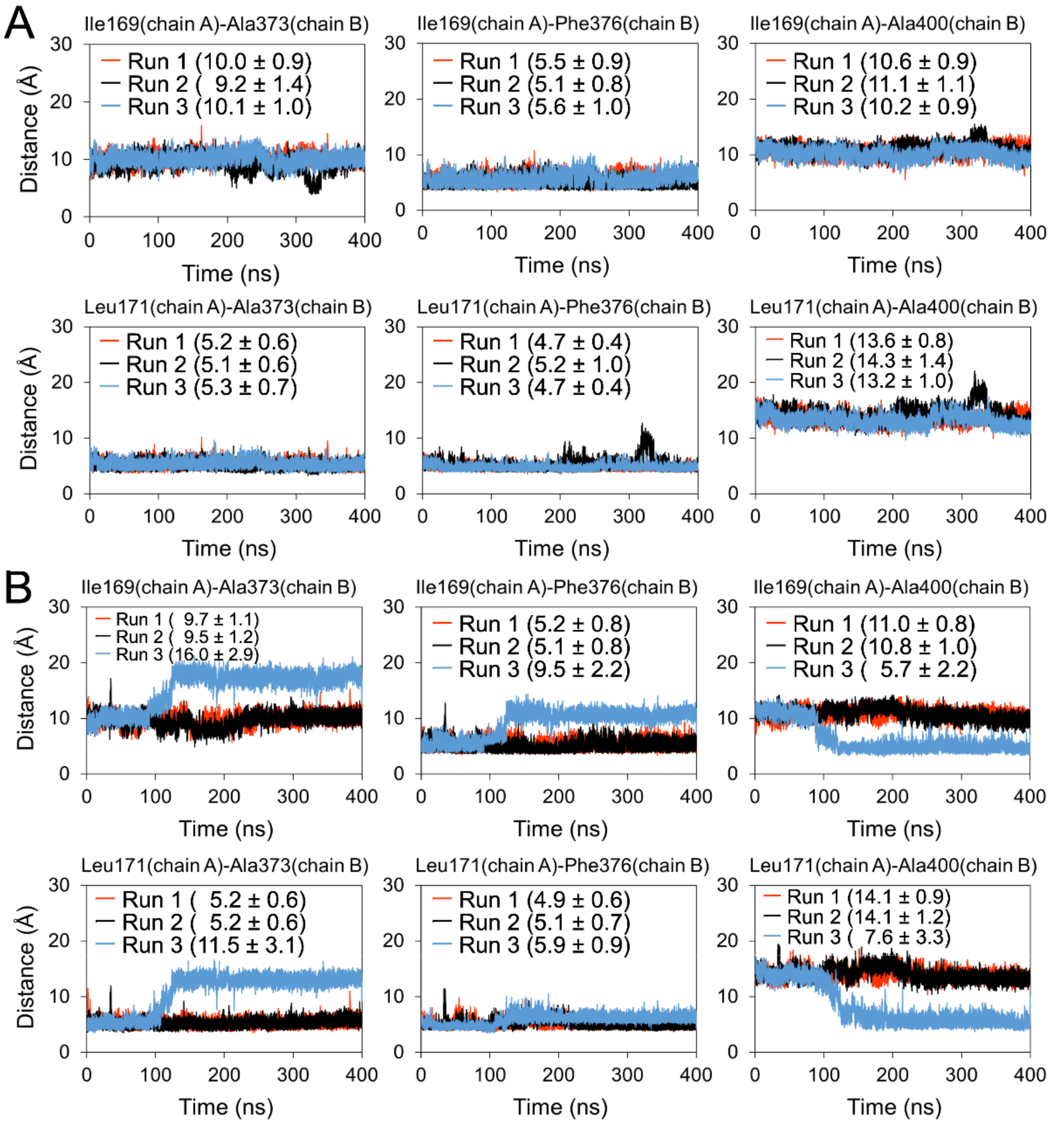


**Figure S3.** Distance between the center of gravity of Ile169 or Leu171 in chain A and Ala373 or Phe376 or Ala400 in chain B during the simulations of the **A.** EC1-4WT homodimer or **B.** EC1-4K115E homodimer. Only Run 3 of the simulations of EC1-4K115E homodimer showed a significant increase or decrease in the middle of the simulations. Averages and standard deviations from 40 to 400 ns of each simulation are shown in the graph in angstrom unit.

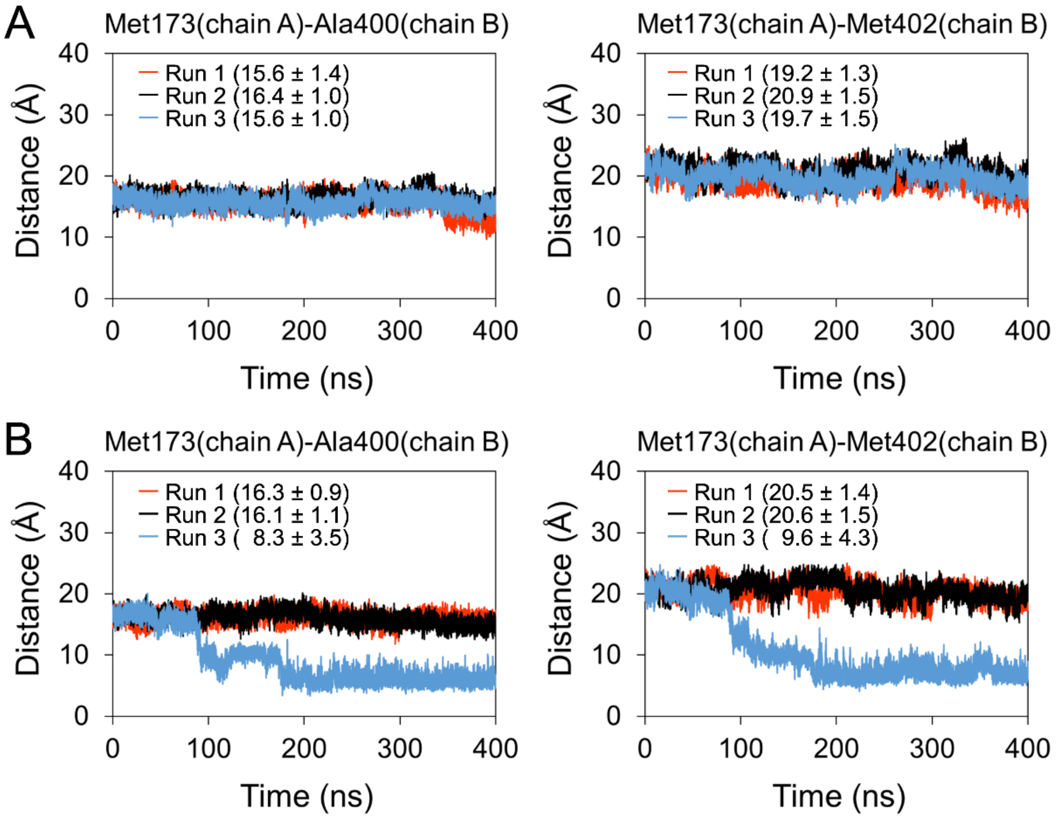


**Figure S4.** Distance between the center of gravity of Met173 in chain A and Ala400 or Met402 in chain B during the simulations of the **A.** EC1-4WT homodimer or **B.** EC1-4K115E homodimer. Only Run 3 of the EC1-4K115E homodimer showed a drastic decrease after approximately 100 ns. Averages and standard deviations from 40 to 400 ns of each simulation are shown in the graph in angstrom unit.

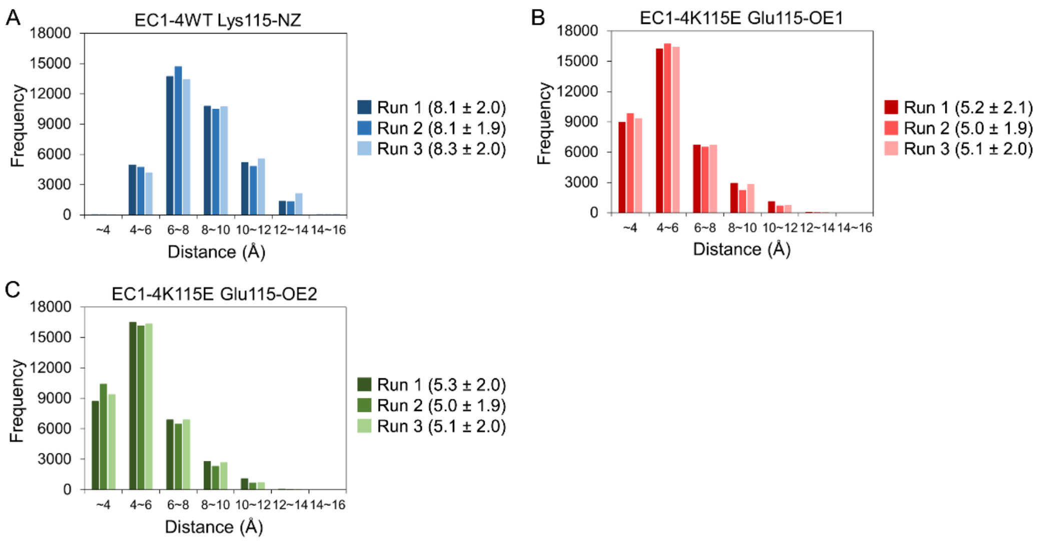


**Figure S5.** Distance between the NZ atom of Lys117 and **A.** NZ atom of Lys115 in the EC1-4WT homodimer, **B.** OE1 atom, or **C.** OE2 atom of Glu115 in the EC1-4K115E homodimer. All atoms belonged to chain A. Averages and standard deviations from 40 to 400 ns of each simulation are shown in the graph in angstrom unit.
